# Stimulus-dependent contrast sensitivity asymmetries around the visual field

**DOI:** 10.1101/2020.06.30.181156

**Authors:** Marc M. Himmelberg, Jonathan Winawer, Marisa Carrasco

**Author notes:** **Corresponding Author:** Marc M. Himmelberg, Department of Psychology, 6 Washington Place, New York University, NY, 10003.

## Abstract

Asymmetries in visual performance at isoeccentric locations are known as *performance fields*. At a fixed eccentricity, visual performance is best along the horizontal, intermediate along the lower vertical, and poorest along the upper vertical meridian. These *performance fields* are pervasive across a range of visual tasks, including those mediated by contrast sensitivity. However, contrast *performance fields* have not been characterized with a systematic manipulation of stimulus spatial frequency, eccentricity, and size; three parameters that constrain contrast sensitivity. Further, individual differences in *performance fields* measurements have not been assessed. Here, we use an orientation discrimination task to characterize the pattern of contrast sensitivity across four isoeccentric locations along the cardinal meridians, and to examine whether and how this asymmetry pattern changes with systematic manipulation of stimulus spatial frequency (4 cpd to 8 cpd), eccentricity (4.5° to 9°), and size (3° visual angle to 6° visual angle). Our data demonstrate that contrast sensitivity is highest along the horizontal, intermediate along the lower vertical, and poorest along the upper vertical meridian. This pattern is consistent across stimulus parameter manipulations, even though they cause profound shifts in contrast sensitivity. Eccentricity-dependent decreases in contrast sensitivity can be compensated for by scaling stimulus size alone. Moreover, we find that individual variability in the strength of *performance field* asymmetries is consistent across conditions. This study is the first to systematically and jointly manipulate, and compare, contrast *performance fields* across spatial frequency, eccentricity, and size, and to address individual variability in *performance fields*.

---

Human visual perception changes as a function of visual field eccentricity and polar angle (Baldwin, Meese, & Baker, 2012; Harrison, 1953; Hering, 1899; Kelly, 1977; Robson & Graham, 1981; Strasburger, Rentschler, & Jüttner, 2011; Traquair, 1928; Wertheim, 1894). At a fixed eccentricity, visual performance is better along the horizontal meridian (HM) when compared to the vertical meridian (VM) – a phenomenon termed the *Horizontal-Vertical Anisotropy* (HVA). Likewise, performance is better along the lower vertical meridian (LVM) than the upper vertical meridian (UVM) – a phenomenon termed the *Vertical-Meridian Asymmetry* (VMA) (Carrasco, Talgar, & Cameron, 2001). Together these asymmetries are called *performance fields* and have been identified for a range of tasks involving contrast sensitivity (Abrams, Nizam, & Carrasco, 2012; Baldwin et al., 2012; Cameron, Tai, & Carrasco, 2002; Levine & McAnany, 2005; Lundh, Lennerstrand, & Derefeldt, 1983; Pointer & Hess, 1989; Regan & Beverley, 1983; Rijsdijk, Kroon, & van der Wildt, 1980; Robson & Graham, 1981; Rovamo & Virsu, 1979; Silva et al., 2008), contrast appearance (Fuller, Rodriguez, & Carrasco, 2008), spatial resolution (Barbot, Xue, & Carrasco, 2020; Carrasco, Williams, & Yeshurun, 2002; Talgar & Carrasco, 2002), temporal information accrual (Carrasco, Giordano, & McElree, 2004), crowding (Fortenbaugh, Silver, & Robertson, 2015; Greenwood, Szinte, Sayim, & Cavanagh, 2017), visual short term memory (Montaser-Kouhsari & Carrasco, 2009), and motion perception (Fuller & Carrasco, 2009).

*Performance fields* are pervasive; they are present binocularly and monocularly (Barbot et al., 2020; Carrasco et al., 2001), across a range of luminance levels (Carrasco et al., 2001), for different stimulus orientations (Baldwin et al., 2012; Carrasco et al., 2001; Corbett & Carrasco, 2011), eccentricities, and spatial frequencies (SFs) (Baldwin et al., 2012; Cameron et al., 2002; Carrasco et al., 2001; Fuller et al., 2008), and correspond to the retinal location of the stimulus (Corbett & Carrasco, 2011). Further, the HVA and VMA are consistent across attention; both exogenous and endogenous attention result in a uniform boost in performance at all isoeccentric locations, preserving the radial asymmetries found in the absence of spatial attention (Abrams et al., 2012; Cameron et al., 2002; Carrasco et al., 2001; Purokayastha, Roberts, & Carrasco, 2020).

*Performance fields* are typically characterized by measuring performance accuracy at isoeccentric locations, with stimulus contrast held at some average threshold. However, performance accuracy at a fixed contrast is only one measure of performance. Contrast sensitivity itself provides an important complementary measure. Psychophysical contrast sensitivity is mediated by the characteristics of both retinal (Curcio & Allen, 1990; Derrington & Lennie, 1984; Shapley, Kaplan, & Soodak, 1981) and cortical cells (Albrecht & Hamilton, 1982; Boynton, Demb, Glover, & Heeger, 1999; DeValois & DeValois, 1991), and as such, can lead to insights about circuit function in the early visual system. To date, no single study of *performance fields* has jointly manipulated stimulus SF, eccentricity, and size – three parameters that are known to impact contrast sensitivity measurements.

As stimulus SF, eccentricity, and size are non-separable determinants of contrast sensitivity, it is important to independently manipulate these parameters in order to attribute changes in contrast sensitivity to a single parameter (Pelli & Bex, 2013). Contrast sensitivity is dependent upon the stimulus SF (Baldwin et al., 2012; Campbell & Robson, 1968; De Valois, Albrecht, & Thorell, 1982; Kelly, 1977; Robson, 1966; Rovamo, Franssila, & Näsänen, 1992) and eccentricity (Baldwin et al., 2012; Hilz & Cavonius, 1974; Jigo & Carrasco, 2020; Pointer & Hess, 1989; Rovamo & Virsu, 1979). Spatial contrast sensitivity is bandpass around the fovea, peaking at ~4 cycles per degree (cpd) (Campbell & Robson, 1968; Kelly, 1977). Sensitivity decreases monotonically with increasing eccentricity, although the extent of this decrease is dependent upon stimulus SF (Hilz & Cavonius, 1974; Koenderink, Bouman, Bueno, & Slappendel, 1978; Wright & Johnston, 1983). The eccentricity-dependent decrease in contrast sensitivity can be quantitatively linked to cortical magnification; there is reduced cortical representation of increasingly peripheral regions of the visual field (Daniel & Whitteridge, 1961; Horton & Hoyt, 1991). Performance on some tasks can be matched at different eccentricities by *M*-scaling the stimulus size such that the spatial extent of the cortical representations are matched (Rovamo & Virsu, 1979; Rovamo, Virsu, & Näsänen, 1978). Further, contrast sensitivity increases with stimulus size due to spatial integration and is known to saturate for larger stimulus sizes, although the critical point at which this saturation occurs is dependent upon stimulus SF (Carlson, 1982; Howell & Hess, 1978; Jarvis & Wathes, 2008; Rovamo, Luntinen, & Näsänen, 1993).

Seminal studies measuring contrast sensitivity across the visual field have shown that eccentricity-gradient changes in contrast sensitivity differ among the cardinal meridians (Pointer & Hess, 1989; Regan & Beverley, 1983; Rijsdijk et al., 1980; Robson & Graham, 1981; Rovamo & Virsu, 1979), and these studies support a trend of contrast sensitivity that resembles the HVA and VMA. Further, contrast response functions have been measured at isoeccentric locations using different SFs (Cameron et al., 2002). These data show that the dynamic range of the contrast response function reflect HVA and VMA, and these asymmetries become exacerbated at higher SFs. These contrast asymmetries progressively weaken with increasing angular distance from the VM, demonstrating that the *performance field* effects are not general hemi- or quarter-field effects, but rather are most pronounced at the meridians (Abrams et al., 2012). Similarly, *performance fields* asymmetries for SF sensitivity also weaken with increasing angular distance from the VM (Barbot et al., 2020). Notably, these studies are yet to measure contrast-defined *performance fields* across different stimulus eccentricities or sizes. Finally, the HVA and VMA are found consistently across observers, however, there is individual variation in the strength of these asymmetries, and this variation has not been thoroughly studied. Given that contrast *performance fields* have been generally established, it is important to assess how and whether they change with the manipulation of parameters known to affect contrast sensitivity within individual observers.

Here, we ask what is the pattern of contrast sensitivity across isoeccentric locations, and does this pattern change with systematic manipulations of stimulus SF, eccentricity, and size? Given their pervasive nature, one might predict that the *performance fields* pattern will be maintained across the shifts in contrast sensitivity that are known to occur from changing these key parameters. Jointly, and in accordance with previous studies, one might predict that the shape of this pattern would become more pronounced with increasing stimulus SF and eccentricity. Further, we ask whether there are reliable differences in the variability of HVA and VMA asymmetry strengths for individual observers, both within and among stimulus conditions. Measurements of asymmetry are important as they are dependent upon an observer’s *difference* in contrast sensitivity at the cardinal meridian, regardless of an observer’s *overall* sensitivity. As HVA and VMA asymmetries are thought to be linked to retinal and cortical organization, we expect individual variability in the size of these asymmetries *within* stimulus conditions. Moreover, we hypothesize that we will find consistency in the relative size of individual observer asymmetries when comparing *among* different stimulus conditions.

To test these hypotheses, we used an orientation discrimination task to measure psychophysical contrast sensitivity at four isoeccentric locations along the cardinal meridians, using four stimulus conditions that varied in stimulus SF, eccentricity, and size. We found that the pattern of contrast sensitivity around the visual field was consistent with the HVA and VMA. This pattern was preserved across these manipulations, despite dramatic shifts in contrast sensitivity at all locations. First, we found that, as expected, increasing the SF or eccentricity of the stimulus decreased contrast sensitivity. Second, the eccentricity-dependent decrease in sensitivity was compensated for by doubling the stimulus size. Third, we found that whereas the *direction* of the HVA and VMA were highly consistent across observers and conditions, the strength of these asymmetries was variable across observers, and the strength of the HVA was variable across conditions.

## Method

### Observers

Nine observers (6 males and 3 females, aged 22-37), including an author (M.M.H), participated in the study and were recruited from New York University. All observers had normal or corrected-to-normal vision. Each observer completed four 1-hour psychophysics sessions. All observers provided informed consent before participating in the study. The experiment was conducted in accordance with the Declaration of Helsinki and was approved by the New York University ethics committee on activities involving human subjects.

### Apparatus and set up

Observers completed each session in a darkened, sound-attenuated room. Stimuli were generated using an Apple iMac (3.2 GHz, Intel Core i3) in MATLAB 2017a (Mathworks, Natick, MA, USA) using the MGL Toolbox (http://gru.stanford.edu/doku.php/mgl/overview). Stimuli were presented on a 21-inch ViewSonic G220fb CRT monitor (1280 x 960 resolution, 100Hz). Gamma correction was performed using a ColorCal MKII colorimeter (Cambridge Research Systems, Rochester, Kent, UK) resulting in a linear relation between frame buffer values and luminance. Observers viewed the monitor display binocularly and their head was stabilized using a chin rest that was positioned 57 cm from the monitor. An Eyelink 1000 eye tracker (SR Research, Ottawa, Ontario, Canada) with a sampling rate of 1000 Hz was used to measure fixation.

### Stimuli

The stimulus consisted of a Gabor patch (a sinusoidal grating embedded in a Gaussian envelope) presented on a uniform grey background. The Gabor patch could be oriented ±15° from vertical. The display contained a central fixation cross (0.5° across) and a set of four stimulus placeholders to eliminate any spatial uncertainty about the potential locations that the Gabor could appear. The placeholders were located at four polar angle locations: 0° at the right horizontal meridian (RHM), 90° at the upper vertical meridian (UVM), 180° at left horizontal meridian (LHM), and 270° at the lower vertical meridian (LVM). The eccentricity and size of the placeholder depended upon stimulus condition. The fixation cross and placeholders remained on the display across the entire experiment.

The Gabor patch differed across the four stimulus conditions. The Gabor was scaled from a ‘baseline condition’ (*condition 1*) by doubling either eccentricity, SF, or eccentricity *and* size. Each experimental session was dedicated to testing one condition as to not introduce uncertainty about the SF or eccentricity of the Gabor. Condition ordering was randomized across sessions. In *condition 1 – baseline* (**Figure 1A**), the Gabor was presented at 4.5° eccentricity, had a SF of 4 cycles per degree (cpd) and covered 3° visual angle (dva) (σ = 0.43°). This condition was chosen as a baseline as the parameters were similar to those used in previous investigations of contrast *performance fields* (Abrams et al., 2012; Cameron et al., 2002). In *condition 2 – SF* (**Figure 1B**), we independently manipulated SF. The Gabor was presented at 4.5° eccentricity and had a SF of 8 cpd, and again covered 3 dva (σ = 0.43°). In *condition 3 - eccentricity* (**Figure 1C**), we independently manipulated eccentricity. The Gabor was presented at 9° eccentricity, had a SF of 4 cpd, and covered 3 dva (σ = 0.43°). To test the effect of stimulus size on any eccentricity-dependent changes in contrast sensitivity, we repeated *condition 3 – eccentricity* and doubled the stimulus size. Thus, in *condition 4 – size and eccentricity* (**Figure 1D**), the Gabor was presented at 9° eccentricity, had a SF of 4 cpd, and was 6 dva (σ = 0.86°) in size. Further, this condition allowed us to isolate the effect of size by comparing between *condition 3 – eccentricity* and *condition 4 – eccentricity and size*, in which the only stimulus change was size.

**Figure 1.**
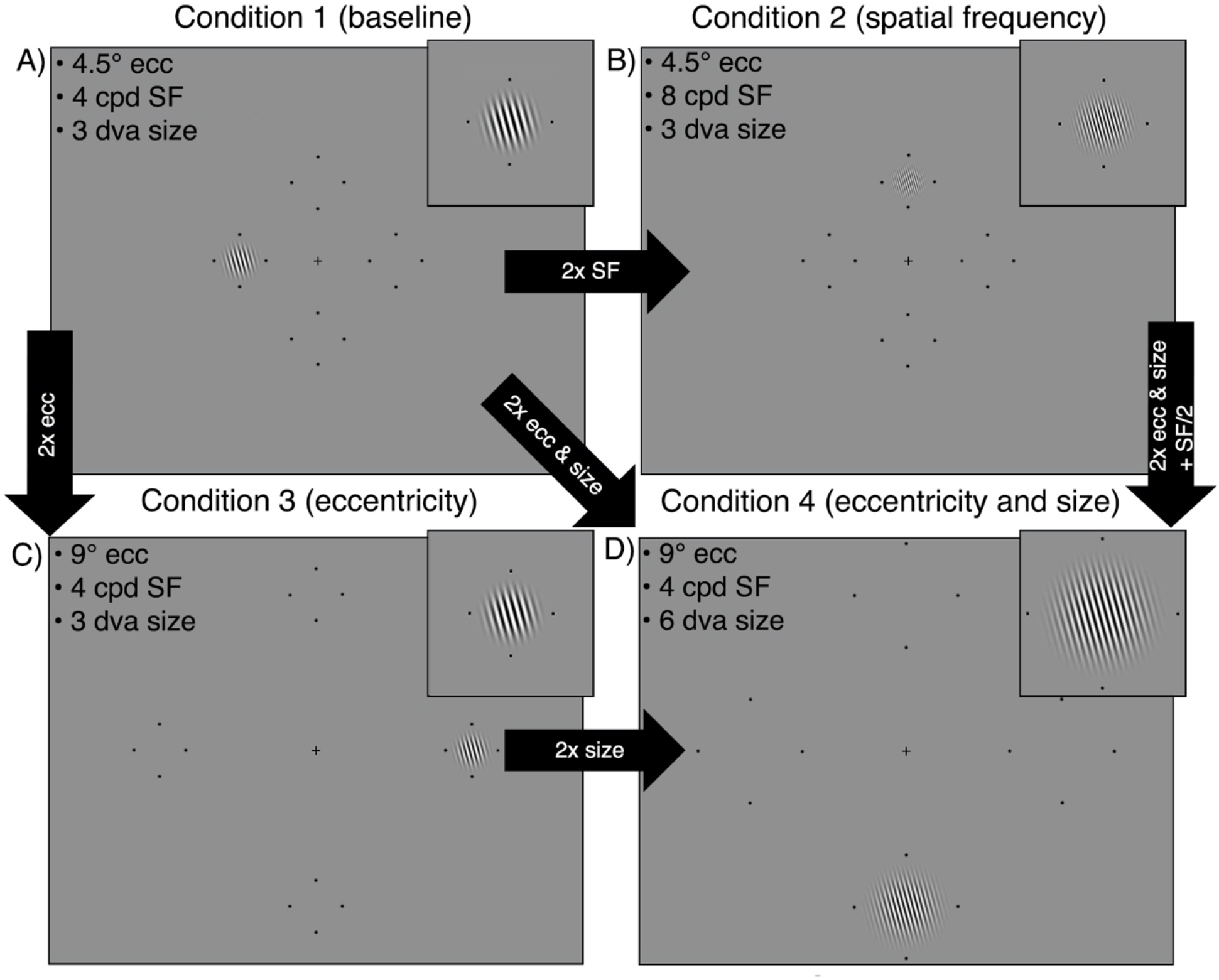
Examples of the four stimulus conditions, each shown at one of the four possible isoeccentric locations. A) *Condition 1* is a ‘baseline condition’ for comparison with the other three conditions. B) In *condition 2* we double the stimulus SF while eccentricity and size are maintained. C) In *condition 3* we double stimulus eccentricity while SF and size are maintained. D) In *condition 4* we double the stimulus eccentricity and size when compared to baseline, while SF is maintained. Placeholders have been enlarged for visibility.

### Experimental Design

Observers completed a two-alternative forced choice (2AFC) orientation discrimination task in which stimulus contrast was titrated using four randomly interleaved 3-down 1-up staircases via parameter estimation by sequential testing (PEST) (Taylor & Creelman, 1967). One staircase was dedicated to each of the visual field locations (RHM, UVM, LHM, LVM). A schematic of a single trial is presented in **Figure 2**. Each trial began with 1000 ms of central fixation (measured using an eye tracker) followed by a 200 ms pre-stimulus cue at fixation to indicate the onset of the trial. This was followed by a 60 ms inter-stimulus-interval (ISI). Next, a Gabor patch was presented at one of the four possible locations for 120 ms. The Gabor was tilted 15° left or right from vertical. There was a 40 ms ISI following the offset of the Gabor patch. This was followed by a 660 ms response cue, a small line extending from the fixation cross. This line indicated the location at which the Gabor patch had appeared to eliminate uncertainty regarding the target location. A brief auditory tone indicated that the observer had 5000 ms to respond, via keyboard press, whether the Gabor was oriented left or right from vertical. Observers were given auditory feedback in the form of a tone to signal whether their response was correct or incorrect. This was followed by a 1000 ms inter-trial-interval (ITI) before the onset of the next trial. Michelson contrast of the Gabor stimulus was titrated across trials in accordance with PEST rules (Michelson, 1927). Trials in which an observer broke fixation (i.e. deviated beyond 1.5 dva from the fixation cross) were aborted and ‘Please fixate’ was presented on the screen. These aborted trials, and any trials in which observers took more than 5 s to respond, were added to the end of the experimental block. Observers were able to break fixation during the response period and during the ITI.

**Figure 2.**
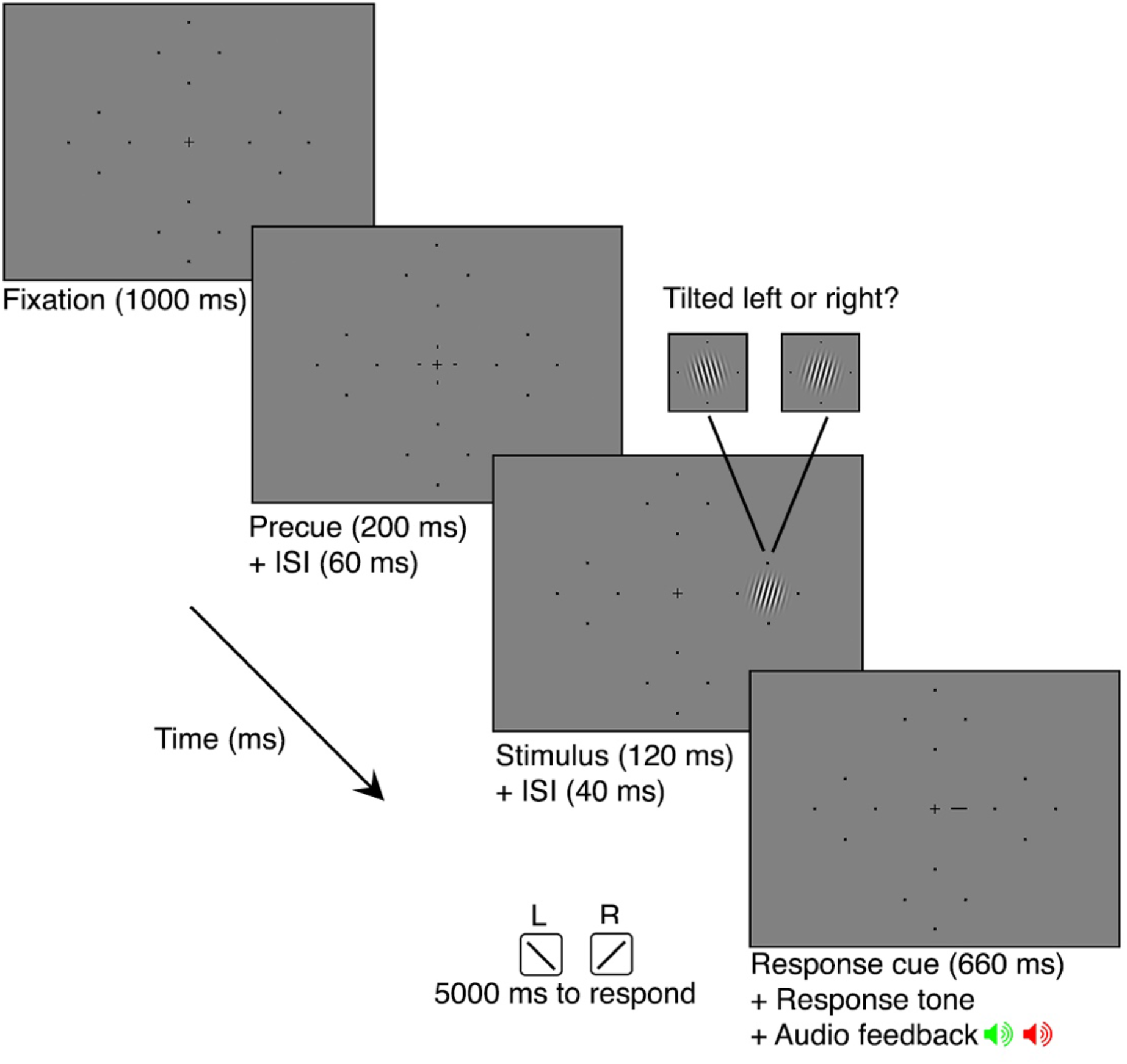
Schematic of a single trial sequence. The Gabor patch could appear in any of the four locations on the screen (denoted by the placeholders) and could be tilted 15° left or right from vertical. Placeholders have been enlarged for visibility.

Each of the four sessions consisted of five independent blocks of 200 trials for a total of 1000 trials per condition, and a total of 250 trials per location for each condition. At the half-way point of each block, observers were given a 20 s break before completing the remaining trials. Observers were given the opportunity to take a break between each block. Before each session, observers completed one block of 24 practice trials (with the Gabor stimuli set to 100% contrast) to ensure that they were comfortable with the task and understood the event timing.

### Analysis

To derive the 75%-correct contrast discrimination threshold for each location and stimulus condition, Weibull functions were fit to each observers data using maximum likelihood estimation (Prins & Kingdom, 2018):

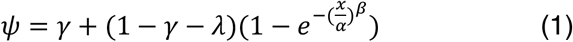

where *ψ* is proportion of correct responses, *x* is the contrast of the stimulus, *γ* is the guessing rate is which was set to 50%, *λ* is the lapse rate which was held constant at 0.01, α is the contrast threshold, and *β* is the slope. Contrast sensitivity values were calculated as the reciprocal of the threshold value. Preliminary analysis found that there was no systematic difference in contrast sensitivity for the LHM and RHM for any of the stimulus conditions (*p* > .05), thus, these data were averaged together in further analysis.

Contrast sensitivity values were entered into two ANOVAs with observer as a random variable; the first tested the HVA and the second tested the VMA. To test for the HVA, we ran a 2 x 4 repeated measures ANOVA to test the effects of visual field location (HM and VM) and stimulus condition (*condition 1* – *baseline, condition 2 – doubling SF, condition 3 – doubling eccentricity*, and *condition 4 – doubling eccentricity and size*) on contrast sensitivity. For the VMA, we ran a 2 x 4 repeated measures ANOVA to test the effects of visual field location (UVM and LVM) and stimulus condition (*condition 1* – *baseline, condition 2 – doubling SF, condition 3 – doubling eccentricity*, and *condition 4 – doubling eccentricity and size*). Mauchly’s test of sphericity was met for all effects and Sidak corrections were applied to post-hoc paired t-tests for all possible comparisons.

## Results

### Contrast sensitivity is dependent upon stimulus parameters and visual field location

The ANOVA to test for the HVA found significant main effects for visual field location (*F*(1,8) = 166.73, *p* < .001, 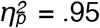) and stimulus condition (*F*(3,24) = 164.97, *p* < .001, 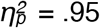), as well as the visual field location by stimulus condition interaction (*F*(3,24) = 119.18, *p* < .001, 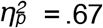). The ANOVA to test for the VMA found significant main effects for visual field location (*F*(1,8) = 30.04, *p* < .01, 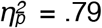) and stimulus condition (*F*(3,24) = 128.59, *p* < .001, 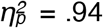, as well as the visual field location by stimulus condition interaction (*F*(3,24) = 7.29, *p* < .01, 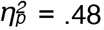). There was no effect of biological sex on contrast sensitivity, nor did biological sex interact with stimulus location or condition (*p* > .1).

### HVA and VMA for contrast sensitivity are consistent across stimulus conditions

We looked at the relevant post-hoc paired t-tests and confirmed that contrast sensitivity changed around the visual field in accordance with the HVA and VMA for each of the four stimulus conditions. The data from *condition 1 – baseline* (**Figure 3A**) demonstrated a clear HVA, with higher contrast sensitivity measurements at the HM than the VM (*p* < .001, mean difference (MD) = 15.46). Likewise, a clear VMA was found with higher contrast sensitivity measurements at the LVM than the UVM (*p* < .05, MD = 5.41). This was also the case for *condition 2 – doubling SF* (**Figure 3B**), HVA (*p* < .001, MD = 2.94) and VMA (*p* < .01, MD = .87), *condition 3 – doubling eccentricity* (**Figure 3C**), HVA (*p* < .001, *MD* = 10.13) and VMA (*p* < .05, MD = 3.08) and *condition 4 – doubling eccentricity and size* (**Figure 3D**), HVA (*p* < .01, MD = 8.67) and VMA (*p* < .01, MD = 5.82).

**Figure 3.**
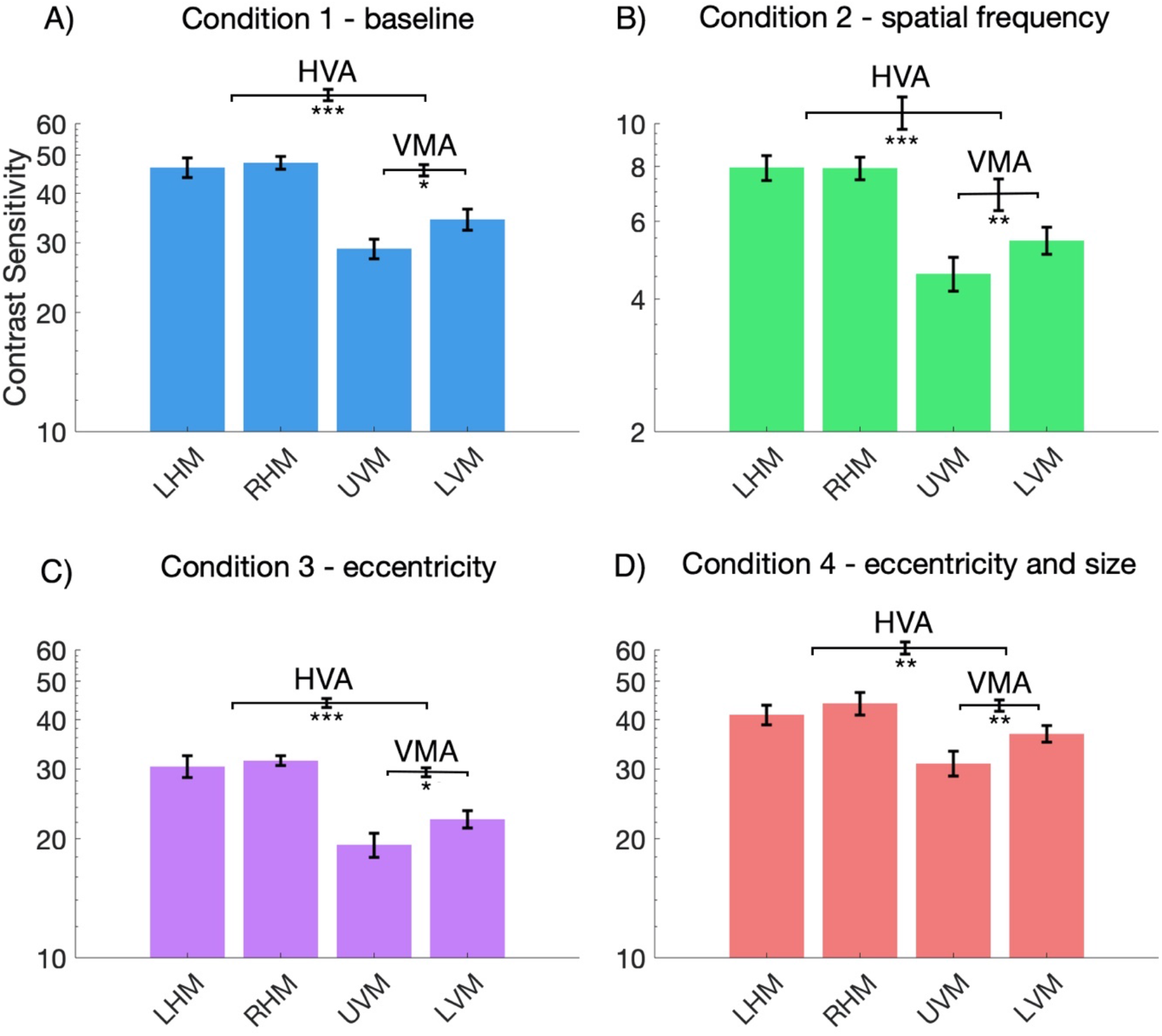
Mean contrast sensitivity plotted as a function of stimulus location for each of the stimulus conditions (n = 9). Note the axis range for panel B) *condition 2 - SF* is different from the other conditions, indicating lower contrast sensitivity for this condition. Error bars are ±1 standard error of the mean (SEM) of each condition (plotted on individual bars) or of the difference between conditions (plotted on the horizontal brackets). * p < .05, ** p < .01, *** p < .001.

To examine variability of the HVA, and the relation between contrast sensitivity on the HM and the VM, contrast sensitivity values for the nine observers were plotted in **Figure 4**. Each data point represents contrast sensitivity values for an individual observer, plotted for the HM (averaged across LHM and RHM) against the VM (averaged across UVM and LVM). The black line represents equal sensitivity between the horizontal and vertical meridians. Generally, contrast sensitivity values fell above the line, indicating that sensitivity was consistently higher along the HM than the VM. Further, we found significant positive correlations between the HM and VM contrast sensitivity measurements for all conditions (*p* < .05), indicating that observers who were more sensitive along the HM were similarly more sensitive along the VM.

**Figure 4.**
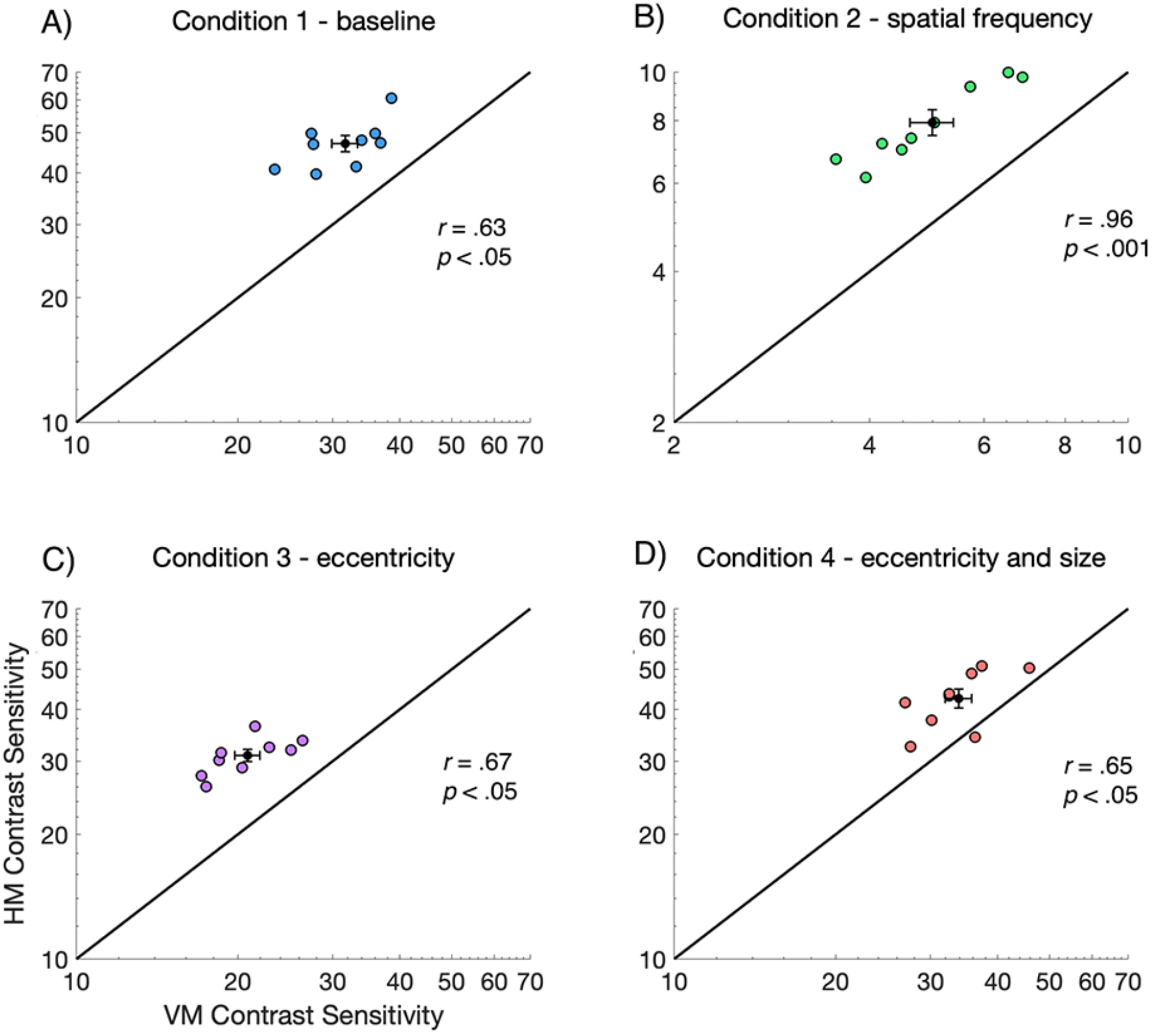
Individual contrast sensitivity data from the horizontal meridian (averaged across left and right) plotted against data from the VM (averaged UVM and LVM) (n=9). The group mean ±1 SEM is presented as the black data point and error bars. Contrast sensitivity is higher along the HM than the VM for all conditions and there is a significant positive correlation between HM and VM contrast sensitivity measurements. Note that the scale of B) *condition 2 - SF* is different from the other conditions, indicating lower contrast sensitivity for this condition. Each data point represents data for a single observer.

Similarly, we examined the variability for the VMA, and the relation between contrast sensitivity on the LVM and the UVM, by plotting LVM versus UVM contrast sensitivity values for the nine observers in **Figure 5**. Each data point represents the contrast sensitivity value for a single observer. Here, data falling above the black line indicates higher sensitivity along the LVM than the UVM. A small subset of observers showed a negligible VMA (i.e. contrast sensitivity was similar for the UVM and LVM). All conditions showed a significant positive correlation between UVM and LVM contrast sensitivity measurements (*p* < .05), indicating that observers who were more sensitive on the UVM were also more sensitive on the LVM.

**Figure 5.**
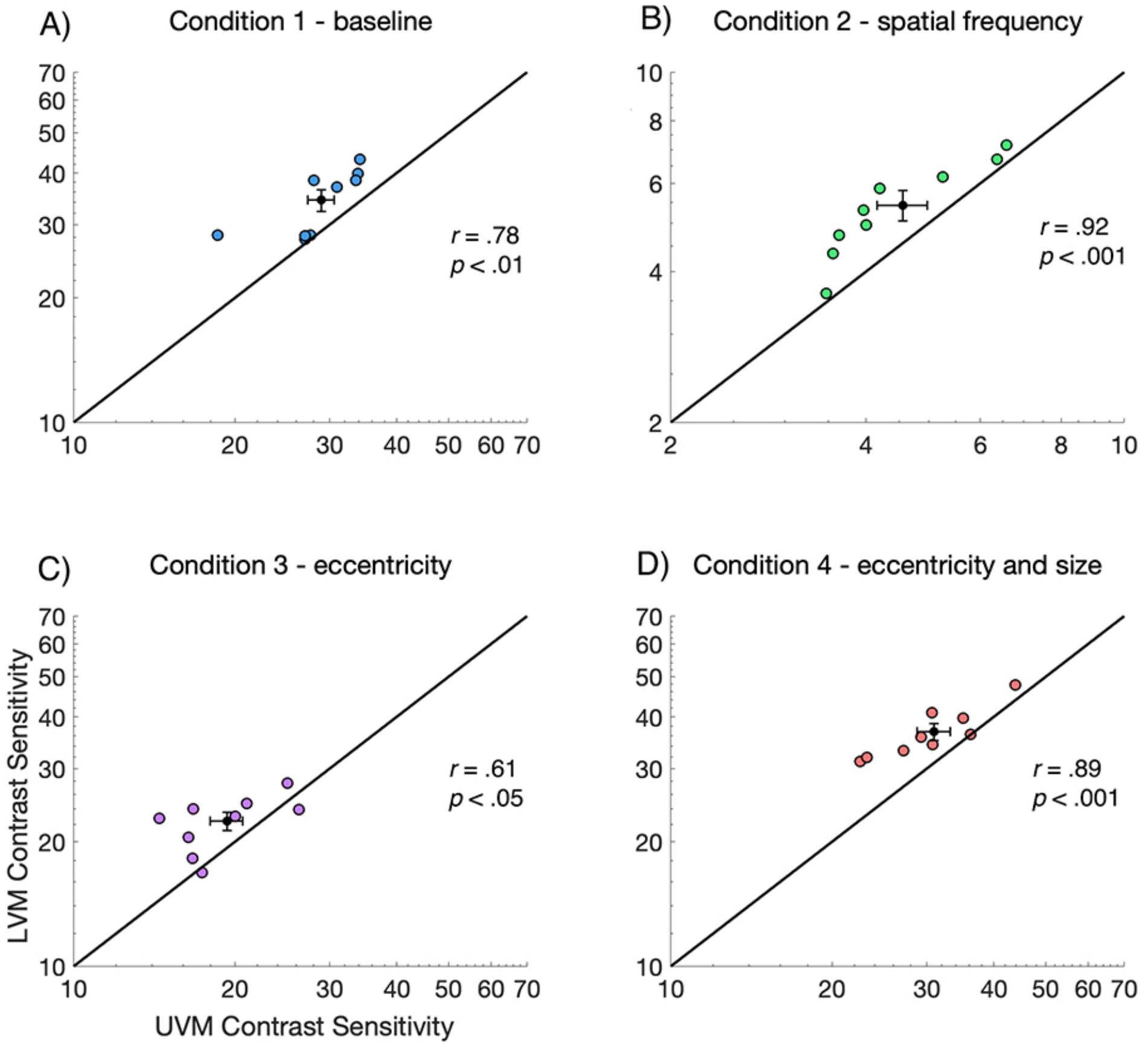
Individual contrast sensitivity data from the UVM plotted against data from the LVM (n=9). The mean value is presented as the black data point and error bars are ±1 SEM. Contrast sensitivity is generally higher along the LVM than the UVM for all conditions and there is a significant positive correlation between UVM and LVM contrast sensitivity measurements. Note that the scale of B) *condition 2 - SF* is different from the other conditions, indicating lower contrast sensitivity for this condition. Each data point represents data for a single observer.

### Contrast sensitivity around the visual field changes as a function of stimulus SF and eccentricity, and doubling stimulus size normalizes contrast sensitivity to baseline

Next, we looked at post-hoc paired t-tests to explore differences in contrast sensitivity among the four stimulus conditions at the different visual field meridians. These analyses revealed that contrast sensitivity was significantly different among stimulus conditions at each of the locations tested (averaged HM, averaged VM, UVM, and LVM), for all possible stimulus condition comparisons (*p* < .01), except when comparing between *condition 1 – baseline* and *condition 4 – doubling eccentricity and size* (*p* > .1).

As illustrated in **Figure 6,** contrast sensitivity was highest at all locations for *condition 1* – *baseline* (blue lines) and *condition 4* – *doubling eccentricity and size* (red lines), with overlap between these two conditions at all four visual field locations. Relative to *condition 1 – baseline* (blue lines), contrast sensitivity decreased across all visual field locations for *condition 3 – doubling eccentricity* (purple lines) and more so for *condition 2 – doubling SF* (green lines). Contrast sensitivity values increased when comparing between *condition 3* – *eccentricity* (purple lines) and *condition 4 – eccentricity and size* (red lines), indicating that doubling stimulus size increased contrast sensitivity at all locations. There was no significant difference between *condition 1 – baseline* (blue lines) and *condition 4 – eccentricity and size* (red lines) at any visual field meridian.

**Figure 6.**
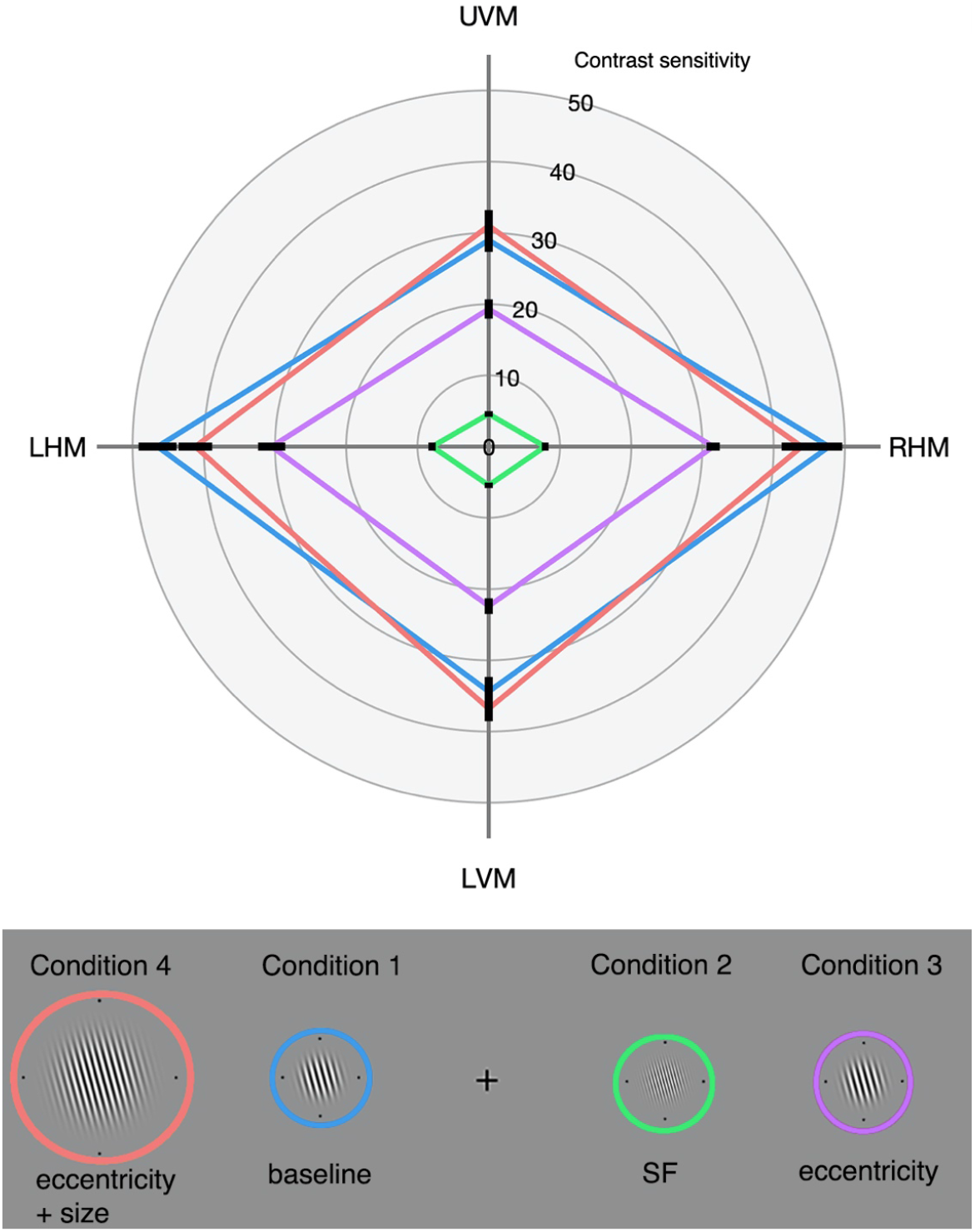
Mean contrast sensitivity plotted as a function visual field location for each stimulus condition (n=9). Contrast measurements were significantly different along each meridian between all conditions, except when comparing between *condition 1* (blue) and *condition 4* (red). Error bars are ±1 SEM.

### HVA strength, but not the VMA strength, changes with stimulus condition at the group level

Next, we explored differences in the strength of the HVA and VMA when comparing between stimulus conditions at the group and the individual level. For each observer, the HVA asymmetry strength value was calculated as percent change in contrast sensitivity from the VM to the HM, whereas the VMA asymmetry strength value was calculated as the percent change in contrast sensitivity from the UVM to the LVM. A positive direction in percent change indicates an asymmetry in line with the HVA or VMA, and a larger percent change indicates a relatively stronger asymmetry. We ran two separate one-way repeated measures ANOVAs to test the effect of the four stimulus conditions on HVA and VMA asymmetry strengths at the group level. Mauchly’s test of sphericity was met for both ANOVAs and Sidak corrections were applied to all post-hoc comparisons.

We found a significant main effect of stimulus condition on HVA strength, *F*(3, 24) = 7.40, *p* < .01, 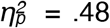. As illustrated in **Figure 7A**, post-hoc t-tests revealed that the HVA asymmetry strength was significantly weaker for *condition 4* – *eccentricity and size* than for *condition 3* – *eccentricity* (*p* < .05) and *condition 2 – SF* (*p* < .01). The HVA asymmetry for *condition 2 – SF* was also marginally stronger than both *conditions 1-baseline* and *3-eccenticity* (both *p* < .1). In contrast, we found no significant effect of stimulus condition on VMA asymmetry strength, *F*(3.24) = 0.53, *p* > .1, 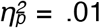, and inspection of **Figure 7B** clearly illustrates the similar strength of the VMA across the four stimulus conditions.

**Figure 7.**
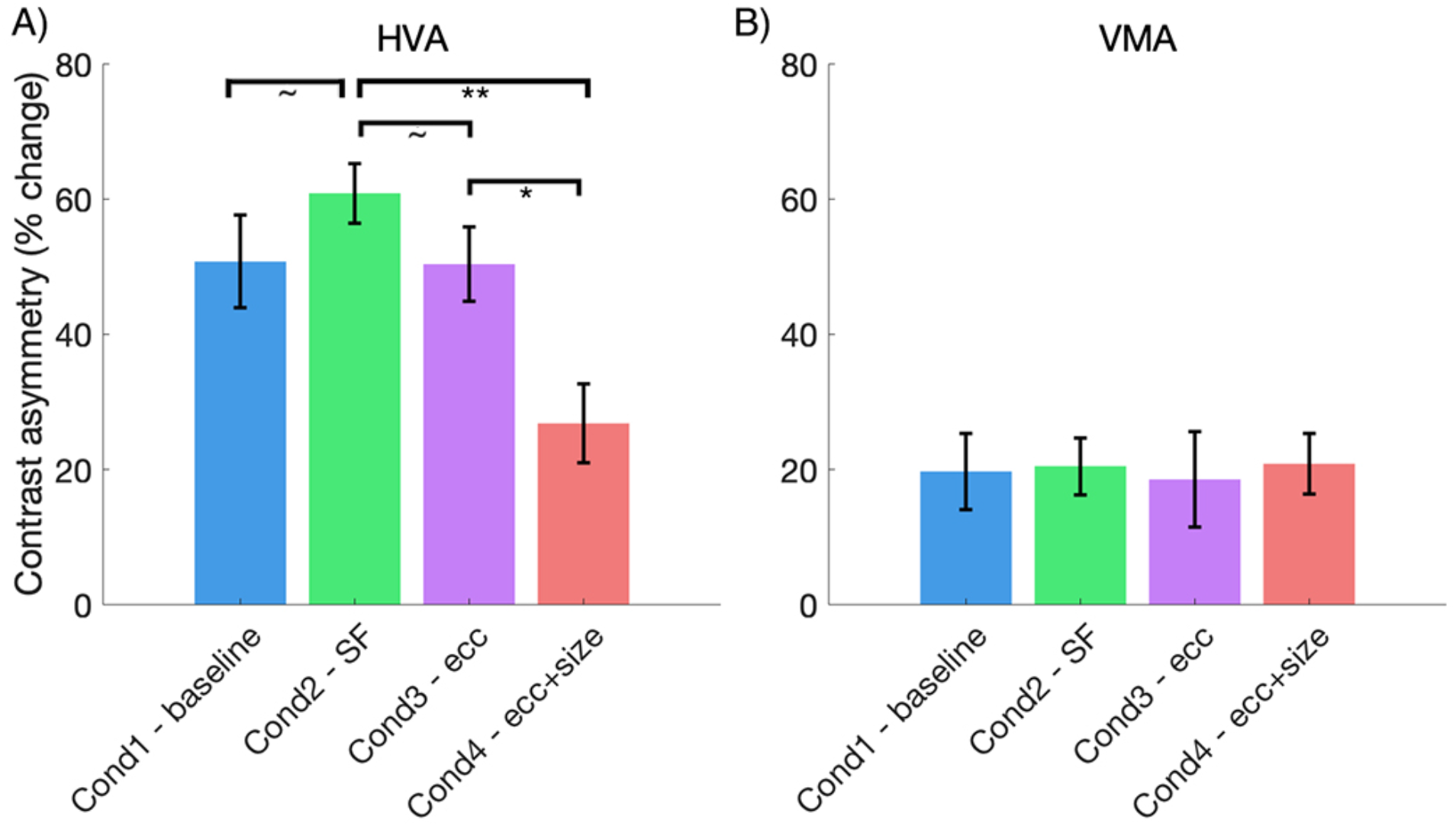
Mean HVA and VMA asymmetry strengths plotted for each stimulus condition (N = 9). HVA asymmetry strength is weaker for *condition 4* – *eccentricity and size* when compared to *condition 2 – spatial frequency* and *condition 3 – eccentricity*, however, VMA asymmetry strength does not change with stimulus condition. Error bars are ±1 SEM. * p < .05, ** p < .01, ~ p < .1.

### Correlations between the strength of the HVA and VMA

Inspection of our data revealed a sizeable spread in the variability of HVA and VMA strength across observers. Thus, we explored the relation between HVA and VMA asymmetry strength for individual observers, at each condition, by plotting observer asymmetry values in **Figure 8**. Data falling above the black line indicated stronger asymmetry for the HVA than for the VMA. We found a significant positive correlation between HVA and the VMA asymmetry strength for *condition 4 – eccentricity and size* (**Figure 8D**; *p* < .05), but no significant correlations for *conditions 1, 2, and 3* (*p* > .1). Additionally, we averaged the HVA and VMA values across all four stimulus conditions and found that there was no significant correlation between the HVA and VMA asymmetry strength (**Figure 8E**; *p* > .1).

**Figure 8.**
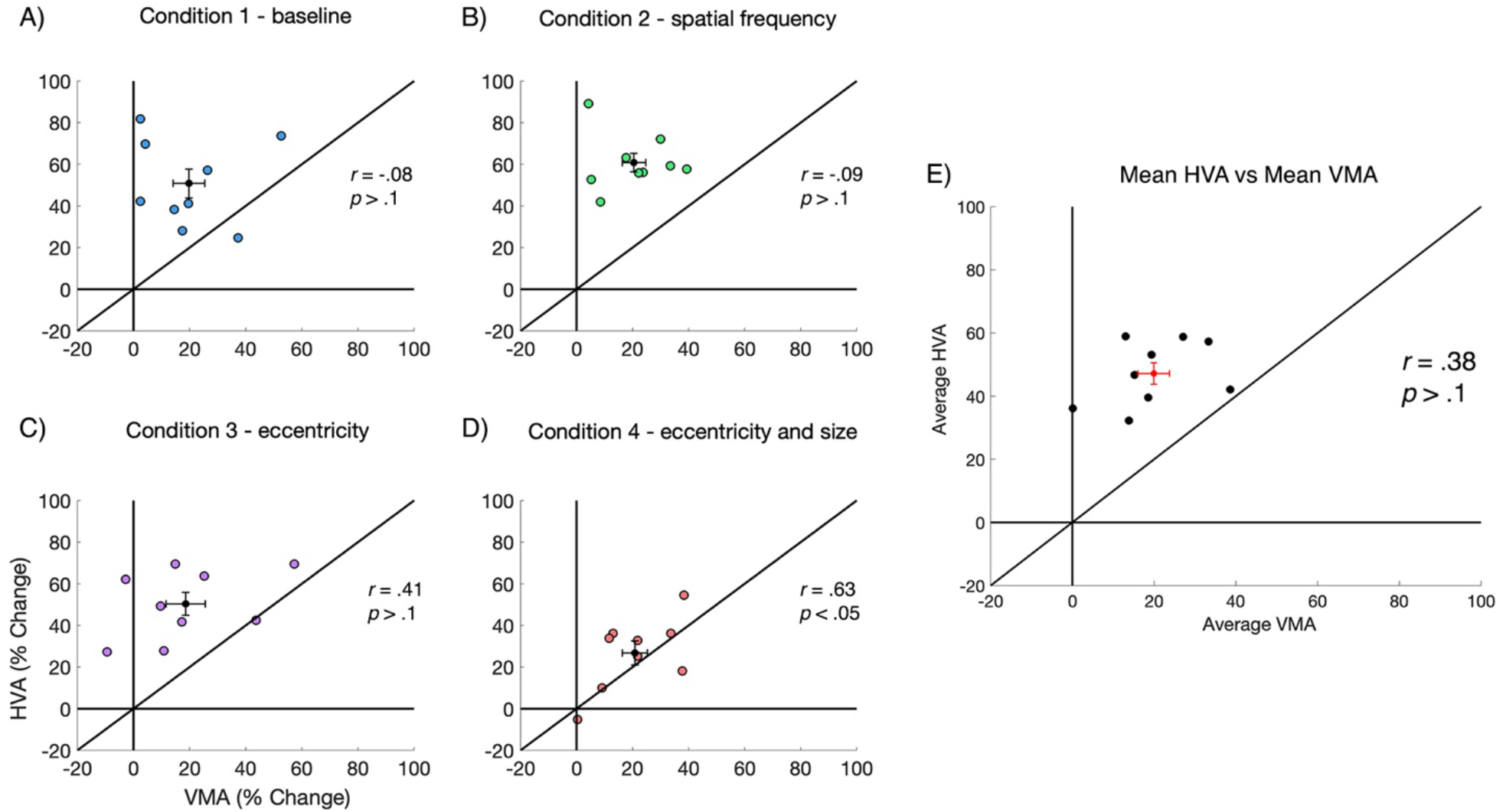
Individual asymmetry data for the HVA strength plotted against VMA strength (n=9). For each condition, the mean value is presented as the black data point and error bars are ±1 SEM. There is no correlation between HVA and VMA strength, except for *condition 4*. Each data point represents data for a single observer.

### Comparing HVA and VMA asymmetry strength among stimulus conditions

To explore how the strength HVA compared among stimulus conditions, we plotted the HVA strength across all possible condition comparisons in **Figure 9**. Data falling above or to the right of the colored lines at ‘0% change’ indicate an asymmetry in line with the HVA, and a larger percent change indicates a relatively stronger asymmetry. If data fall on the black diagonal line it suggests that the asymmetry strength is identical between the two plotted conditions; if the value is above the diagonal line the asymmetry is stronger for the condition plotted on the y-axis, and if it is below it is stronger for the condition plotted on the x-axis.

**Figure 9.**
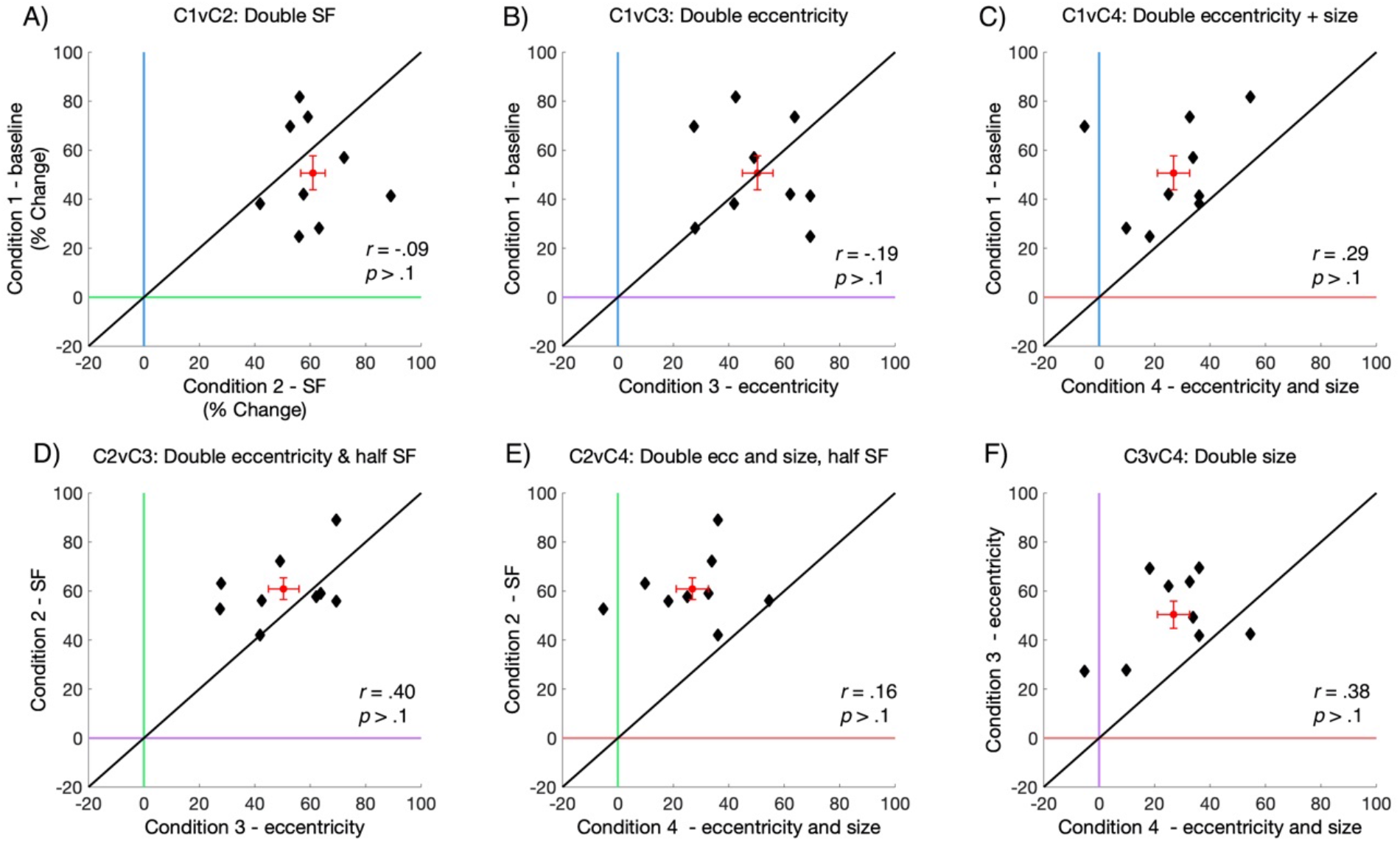
HVA asymmetry strength values plotted for all possible condition comparisons. Each data point representing data from an individual observer (n=9). The red data point represents the mean value with error bars ±1 SEM. Data on the diagonal line indicates an identical HVA asymmetry strength between the two plotted conditions.

We found no significant correlations in HVA strength when comparing across all conditions (*p* > .1), although there was, generally, a positive trend in the HVA strength between conditions. Most notably, these data show that there is substantial variability in asymmetry strength across observers. The bulk of HVA asymmetry values fell between ~40 - 80% in *conditions 1, 2, and 3*, whereas the HVA asymmetry values in *condition 4 – doubling eccentricity and size* were relatively lower, sitting between ~10 – 40%. This indicates that scaling stimulus size resulted in some weakening of the HVA asymmetry.

As there was substantial variability across individual HVA strengths, we further inspected the data in **Figure 9** to make inferences about how the strength of individual HVA asymmetries compared after isolating the effects of SF, eccentricity, and size on contrast sensitivity. Specifically, we found that doubling the stimulus SF from 4 to 8 cpd (**Figure 9A**) caused the HVA asymmetry to become slightly stronger. There was no relative difference in HVA asymmetry strength after doubling stimulus eccentricity from 4.5° to 9° (**Figure 9B**); some observers had higher asymmetry measurements for the baseline condition, but others had higher asymmetry for the doubled eccentricity condition. The HVA was weaker after doubling stimulus eccentricity and size (**Figure 9C**; i.e. cortically magnifying the Gabor). Next, doubling stimulus eccentricity and halving the SF (**Figure 9D**) weakened the strength of the HVA asymmetry, whereas doubling the stimulus eccentricity and the size while halving the SF (**Figure 9E**) also weakened the strength of the HVA asymmetry. Finally, doubling the stimulus size also weakened the strength of the HVA asymmetry (**Figure 9F**).

Similarly, we plotted we plotted the VMA asymmetry strength across all possible condition comparisons in **Figure 10**. Again, data falling above or to the right of the colored lines at ‘0% change’ indicate an asymmetry in line with the VMA, and a larger percent change indicates a relatively stronger asymmetry. If a data point is above the diagonal line, the VMA is stronger for the condition plotted on the y-axis, and if it is below the line the VMA is stronger for condition plotted the x-axis.

**Figure 10.**
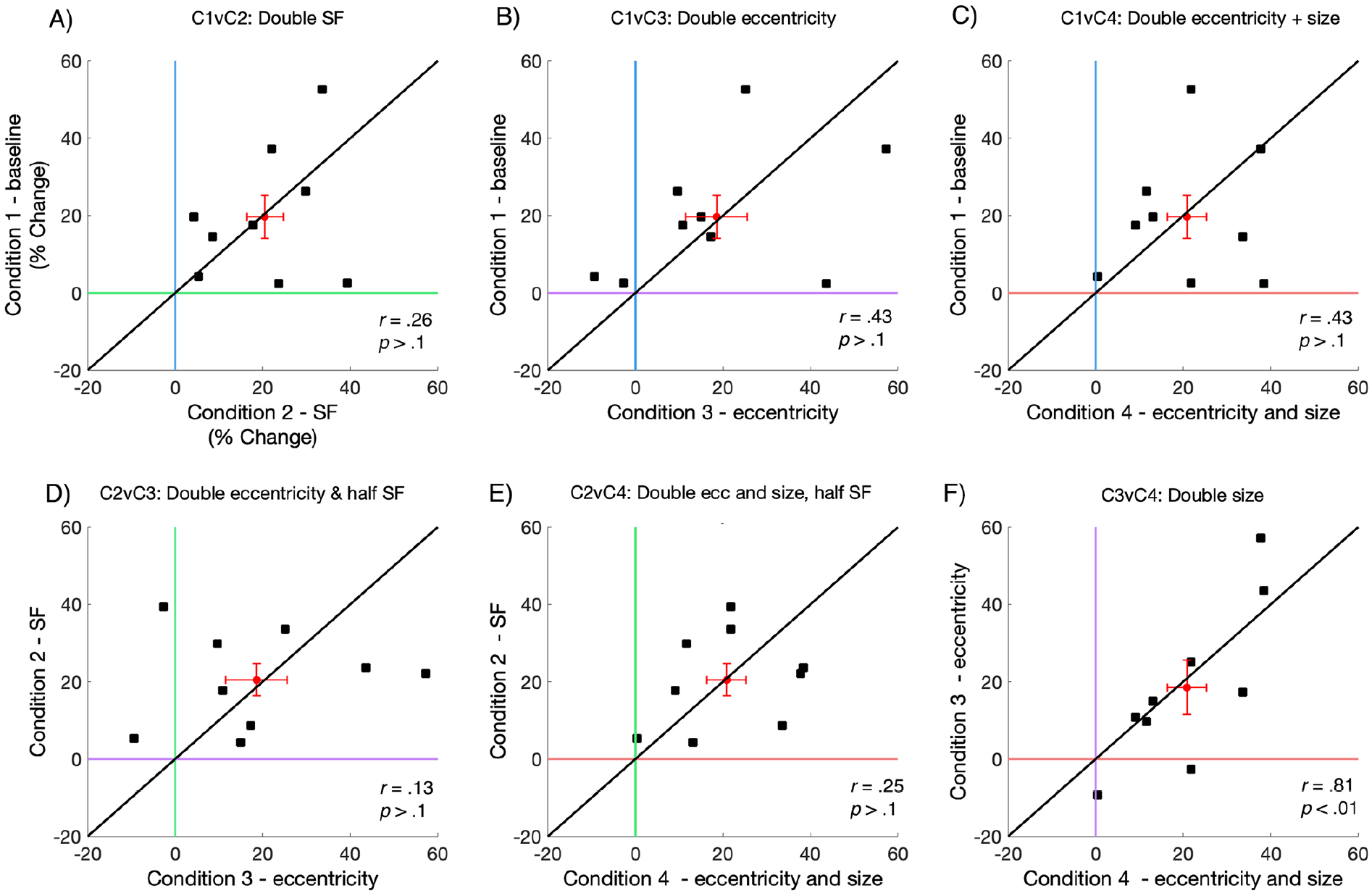
VMA asymmetry strength values plotted for all possible condition comparisons. Each data point representing data from an individual observer (N=9). The red data point represents the mean value with error bars ±1 SEM. Data falling on the diagonal line indicates an identical VMA asymmetry strength between two plotted conditions.

The VMA asymmetry strength was lower than HVA asymmetry strength, with the bulk of VMA values falling between ~10-40% across all conditions. Further, we found no significant correlations for the strength of the VMA strength when comparing between conditions (*p* > .1), except for when comparing between *condition 3 – size* and *condition 4 – size and eccentricity* (**Figure 10F**; *p* < .01). This suggests that at the individual observers VMA strength is similar after doubling the stimulus size. Similar to our observations of VMA strength at the group level (**Figure 7B**), this analysis indicates no reliable differences in the strength of the VMA at the individual level, with data falling both above and below the black diagonal line.

## Discussion

We have characterized contrast *performance fields* across manipulations of stimulus SF, eccentricity, and size – three key parameters that constrain contrast sensitivity. Our measurements show that the pattern of contrast sensitivity at isoeccentric locations is consistent with the HVA and VMA. This pattern is stable after systematically manipulating these key stimulus parameters, even though this provokes dramatic shifts in contrast sensitivity across visual field locations. Further, our data show that eccentricity-dependent decreases in contrast sensitivity can be accounted for by doubling the stimulus size. Finally, we have explored the variability of the HVA and VMA asymmetry strengths at the individual observer level and found consistent differences in these asymmetries across stimulus conditions.

### Contrast performance fields are preserved over manipulations of stimulus spatial frequency, eccentricity, and size

Our data show a clear HVA and VMA for contrast sensitivity. Consistent with previous studies, observers had greater contrast sensitivity on the HM than the VM, and on the LVM than the UVM (e.g. Abrams et al., 2012; Baldwin et al., 2012; Cameron et al., 2002). Thus, observers require more contrast to perform at 75% accuracy at the VM when compared to the HM, and at the UVM when compared to the LVM. These asymmetries were prevalent across systematic manipulations of stimulus SF, eccentricity, and size, even though these manipulations resulted in relatively large shifts in contrast sensitivity.

*Performance field* asymmetries are now known to exist for both contrast sensitivity and spatial resolution – two parameters that underlie the processing of all visual stimuli (Abrams et al., 2012; Barbot et al., 2020; Cameron et al., 2002). This is important, as asymmetries for these low-level mechanisms will impact performance for the higher-order visual processes that dictate how we perceive, and subsequently interact, with our visual environment. Overall, our data are the first to show that contrast *performance fields* are maintained across systematic manipulations of SF, eccentricity, and size, and provide continued evidence for the ubiquitous nature of *performance fields* asymmetries.

### Contrast sensitivity changes with stimulus spatial frequency and eccentricity at all isoeccentric locations

Increasing the SF or eccentricity of the stimulus significantly reduced contrast sensitivity, and this was common across all four meridians. It is well documented that contrast sensitivity is dependent upon stimulus SF and eccentricity (Campbell & Robson, 1968; De Valois et al., 1982; Hilz & Cavonius, 1974; Jigo & Carrasco, 2020; Kelly, 1977; Pointer & Hess, 1989; Robson, 1966; Rovamo et al., 1992; Virsu & Rovamo, 1979). In the baseline condition (4 cpd), contrast sensitivity values fell around ~50 on the HM and ~33 on the VM (corresponding to thresholds at ~2% and ~3%, respectively). Doubling the SF of the Gabor stimulus to 8 cpd profoundly reduced contrast sensitivity; it fell between ~8 on the HM and ~ 4.5 on the VM (corresponding to thresholds at ~12% and ~22%, respectively). Previous data shows that increasing stimulus SF increases the magnitude of the asymmetries (Cameron et al., 2002; Carrasco et al., 2001). After inspecting individual differences in the strength HVA asymmetry, we found that that doubling SF did result in a stronger HVA asymmetry, although this was not significant at the group level, but there was no change in the strength of the VMA. *Performance fields* are typically characterized by measuring performance accuracy at isoeccentric locations, with contrast held at the average threshold of from these locations. Thus, the dramatic difference in contrast sensitivity between the HM and VM (for a high SF stimulus) would suggest that averaging these thresholds together could skew the average threshold, resulting in near perfect performance on the HM and heavily reduced performance on the UVM. This is important for future studies that might aim to investigate the role of SF in *performance fields*.

Consistent with previous work (Virsu & Rovamo, 1979), doubling stimulus eccentricity from 4.5° to 9° resulted in a ~2-fold decrease in contrast sensitivity. This decrease was similar at each of the four locations so that increasing eccentricity did not cause the HVA or VMA to become more pronounced at the individual or the group level. Previous findings indicate that the HVA and VMA become more pronounced at farther eccentricities *and* at higher SFs; this is because the spatial resolution of the visual system decreases with increasing eccentricity (Aghajari, Vinke, & Ling, 2020; Carrasco et al., 2001; Robson & Graham, 1981). Thus, one might expect to find a more pronounced HVA and VMA at 9° eccentricity if we had jointly increased the stimulus SF. Moreover, contrast sensitivity declines more rapidly with eccentricity at more central visual field locations (Pointer & Hess, 1989; Rijsdijk et al., 1980; Rovamo, 1983). Our measurements are taken at parafoveal to near-peripheral eccentricities (4.5° and 9°), and it is possible that larger differences in contrast sensitivity would occur if we compared contrast measurements at more central eccentricities, where the rate of contrast sensitivity change is steepest.

In accordance with previous literature (Carlson, 1982; Howell & Hess, 1978; Jarvis & Wathes, 2008; Rovamo et al., 1993), we found doubling stimulus size (3 dva to 6 dva, at 9° eccentricity) increased contrast sensitivity ~1.5 fold and this effect was found at all isoeccentric locations. The increase in contrast sensitivity with stimulus size is most likely due to spatial integration, in which the visual system summates information across contiguous regions of the visual field to increase sensitivity (Rovamo et al., 1993). Doubling the stimulus size *reduced* the strength of the HVA asymmetry, and inspection of our data showed that this is because there was a relatively large increase in sensitivity for the LVM relative to the HM, which caused an overall weakening of the HVA. Thus, it appears as though increasing stimulus size had a larger effect on contrast sensitivity at the LVM when compared to the other visual field locations.

### Eccentricity-dependent decreases in contrast sensitivity at isoeccentric locations can be accounted for by doubling stimulus size

Our data show that eccentricity-dependent changes in contrast sensitivity could be reversed by doubling the Gabor size without changing the stimulus SF content. This effect was consistent for all four visual field locations. This finding can be interpreted in light of cortical magnification. The amount of cortical space dedicated to processing visual information decreases with increasing visual field eccentricity (Daniel & Whitteridge, 1961; Horton & Hoyt, 1991). Stimulus size can be scaled by some *M*-factor to equalize its cortical representation, consequently equalizing contrast sensitivity measurements taken at different eccentricities (Rovamo, 1983; Rovamo et al., 1978; Virsu & Rovamo, 1979). Although we have not used a precise *M*-factor scaling, a simple geometric doubling of stimulus size accounted for the eccentricity-dependent changes in contrast sensitivity that we observed at all four locations.

To match stimulus representations as closely as possible across different eccentricities, one option is to decrease SF in proportion to the increase in size. The logic is that in cortex, greater eccentricity is associated with both decreased cortical area and tuning to lower SFs (De Valois et al., 1982; Xu, Anderson, & Casagrande, 2007). In our study, the eccentricity-dependent changes in contrast sensitivity were compensated for by scaling stimulus size alone (at least for the intermediate SF range we have tested here). It is possible that eccentricity-dependent changes in visual sensitivity are predominately due to differences in anatomical space (cortical magnification), rather than differences in SF sensitivity that occur as a function of eccentricity. This finds support in previous studies in which magnifying stimulus size alone accounted for eccentricity effects in visual search tasks (Carrasco & Frieder, 1997; Chan & Courtney, 1998), temporal contrast sensitivity (Himmelberg & Wade, 2019), and letter recognition (Anstis, 1974; Farrell & Desmarais, 1990; Higgins, Arditi, & Knoblauch, 1996). If these eccentricity-dependent differences in sensitivity can be accounted for by stimulus size alone, it is reasonable to theorize that isoeccentric differences in contrast sensitivity are also rooted in the differences in the spatial scale of cortical representations (i.e. differences in cortical magnification along the different visual field meridians). If so, the difference in performance could be eliminated by a similar stimulus ‘isoeccentric scaling’.

### Observer HVA and VMA contrast asymmetries are similar for different stimulus conditions

Overall, our data showed significant positive correlations for individual measurements of contrast sensitivity taken at different meridians. Observers who had higher contrast sensitivity along the HM also had higher contrast sensitivity along the VM, and this was the case for all four conditions. These correlations were even stronger for the VMA – those who were more sensitive on the LVM were also more sensitive on the UVM.

Testing the same observers across four conditions in which we systematically varied SF, eccentricity, and size allowed us to inspect individual observer variability in the strength of HVA and VMA asymmetries both within and between stimulus conditions. These measurements rely upon the percent change in contrast sensitivity when comparing between two meridians. The strength of these asymmetries was surprisingly broad across the nine observers. The HVA strength ranged between a ~40% to 80% increase in contrast sensitivity from the VM to the HM (except for *condition 4 – double eccentricity and size*, where the bulk of HVA measurements ranged from 20% to 60%). Comparatively, the VMA was weaker and the bulk of measurements ranked from 10% to 40% change in contrast sensitivity from the LVM to the UVM. At the group level, there was a consistent mean ~20% increase in contrast sensitivity from the UVM to the LVM for all four stimulus conditions.

We also explored individual HVA and VMA asymmetry values when comparing across our different stimulus conditions, which effectively isolated effects of SF, eccentricity, and size. Although the strength of the HVA or VMA did not significantly correlate between condition comparisons (except one; the VMA after doubling stimulus size), we identified a general positive slope in our data; observers with a relatively stronger HVA or VMA asymmetry in one condition tended to have a stronger HVA or VMA asymmetry in another condition. Some consistency in the strength of *performance fields* asymmetries across stimulus conditions might suggest that there is meaningful variability *between* observers, even when this variability is surprisingly large. This variability in *performance field* asymmetries will prove to be informative in exploring their neural substrates.

Following this, we explored how individual trends in the strength of HVA and VMA asymmetries changed after isolating out the effects of manipulating SF, eccentricity, and size. The strength of the HVA asymmetry was weakened as a result of doubling stimulus size alone, by doubling eccentricity and size (and this was similar when SF was halved), and doubling eccentricity and halving the SF. Conversely, we found that HVA asymmetries were strengthened by doubling the stimulus SF for most, but not all, observers. In contrast, changing stimulus SF, eccentricity, or size did not have any visible effect on the strength of the VMA asymmetry. Thus, the VMA (when defined using contrast sensitivity measurements) does not seem to be particularly sensitive to changes in any of these parameters.

Finally, we identified a significant correlation between the HVA and VMA for only one of our four stimulus conditions, *condition 4 – eccentricity and size*. This suggests that the HVA and VMA strengths are largely independent from each other; an observer with a strong HVA asymmetry will not necessarily have a strong VMA asymmetry. If HVA and VMA strengths are independent from each other, one might hypothesize that these two asymmetries also emerge independently from each other across development. A recent report suggests this is indeed the case (Myers & Carrasco, 2020).

### Neural substrates of performance fields asymmetries

Overall, these results demonstrate that *performance fields* asymmetries are pervasive across manipulations of stimulus eccentricity, SF, and size that result in dramatic shifts in contrast sensitivity. The prevalent nature of the HVA and VMA might be due to similar asymmetries across multiple stages of the visual pathway that become progressively more reflective of behavioural measurements as signals travel from retina to cortex. These asymmetries begin at the front-end of the visual system; there is a ~30% increase in retinal cone density between the VM and the HM (Curcio, Sloan, Kalina, & Hendrickson, 1990; Song, Chui, Zhong, Elsner, & Burns, 2011). This asymmetry is amplified at the RGC cells, where there is a ~40% increase in RGCs density between the VM and the HM (Curcio & Allen, 1990; Watson, 2014). However, the recent implementation of a computational observer model has identified that asymmetries in cone sampling and optics can only account for a small portion of isoeccentric differences in contrast sensitivity. Specifically, the model found that a ~30% decrease in contrast sensitivity between the HM and UVM would require a 500% decrease in cone density, which is well beyond the actual density change (Kupers, Carrasco, & Winawer, 2019). Notably, cone densities can vary up by to a factor of 3 across individuals (Curcio, Sloan, Packer, Hendrickson, & Kalina, 1987). If retinal asymmetries can only account for a small fraction of isoeccentric differences in contrast sensitivity, perhaps these differences are exacerbated further downstream.

Cortical magnification in primary visual cortex (V1) decreases as a function of eccentricity (Daniel & Whitteridge, 1961; Engel et al., 1994; Holmes, 1919; Horton & Hoyt, 1991; Inouye, 1909) and has more recently been described to vary as a function of polar angle. fMRI studies have identified a cortical HVA; there is more cortical space dedicated to processing visual information along the HM of the visual field when compared to the VM (Benson et al., 2012; Benson et al., 2019; Silva et al., 2018). Further, there is evidence for a cortical VMA; there is more cortical space dedicated to the LVM than the UVM and this difference decreases gradually towards the HM (Benson et al., 2019; Benson, Kupers, Carrasco, & Winawer, 2020), and there is a 40% increase in BOLD amplitude at the LVM than at the UVM (Liu, Heeger, & Carrasco, 2006). Although these cortical asymmetries are well-described, we do not know the extent to which they can account for the HVA and VMA, or how far they extend beyond early visual cortex. Similar to retinal cone densities, cortical space in V1 can differ by a factor of 3 among individuals (Andrews, Halpern, & Purves, 1997; Dougherty et al., 2003; Stensaas, Eddington, & Dobelle, 1974). The variability in HVA and VMA strength described in the current study could, potentially, be accounted for by similar variability in cortical HVA and VMA asymmetries. We are currently developing computational models to assess whether, and to what extent, different stages of the visual pathway can account for observed behavioral differences in the current and previous studies.

## Conclusion

We have characterized contrast *performance fields* across manipulations of fundamental stimulus parameters that contrast sensitivity measurements are contingent upon. Our data showed that the pattern of contrast sensitivity around the visual field followed the HVA and VMA, and this was consistent after doubling stimulus SF, eccentricity, and size. Doubling stimulus SF and eccentricity caused reductions in contrast sensitivity at all isoeccentric locations, whereas doubling stimulus size at a single eccentricity resulted in increased contrast sensitivity. Further, eccentricity-dependent changes in contrast sensitivity could be accounted for by stimulus size alone, a finding we attribute to cortical magnification. Finally, our data showed general similarities in the strength of individual observer HVA and VMA asymmetries when comparing among stimulus conditions, even though the strengths of these asymmetries were relatively broad at the group level. We hypothesize that the overall breadth of these asymmetries might be linked to individual differences corresponding to neural asymmetries that become progressively more pronounced along the visual pathway, culminating at the cortex. Overall, these findings contribute to the fundamental characterization of how contrast sensitivity changes around the visual field and should be taken into account in models of vision.

## Acknowledgements

NIH-NEI RO1-EY027401 to MC and JW. We would like to thank Antoine Barbot, Antonio Fernández, Michael Jigo, Eline Kupers and Shutian Xue and for their helpful comments on the manuscript.

## Commercial relationships

none

